# Core PCP mutations affect short time mechanical properties but not tissue morphogenesis in the *Drosophila* pupal wing

**DOI:** 10.1101/2022.12.09.519799

**Authors:** Romina Piscitello-Gómez, Franz S Gruber, Abhijeet Krishna, Charlie Duclut, Carl D Modes, Marko Popović, Frank Jülicher, Natalie A Dye, Suzanne Eaton

## Abstract

How morphogenetic movements are robustly coordinated in space and time is a fundamental open question in biology. We study this question using the wing of *Drosophila melanogaster*, an epithelial tissue that undergoes large-scale tissue flows during pupal stages. We showed previously (Etournay et al., 2015) that pupal wing morphogenesis involves both cellular behaviors that allow relaxation of mechanical tissue stress, as well as cellular behaviors that appear to be actively patterned. The core planar cell polarity (PCP) pathway influences morphogenetic cell movements in many other contexts, which suggests that it could globally pattern active cellular behaviors during pupal wing morphogenesis. We show here, however, that this is not the case: there is no significant phenotype on the cellular dynamics underlying pupal morphogenesis in mutants of core PCP. Furthermore, using laser ablation experiments, coupled with a rheological model to describe the dynamics of the response to laser ablation, we conclude that while core PCP mutations affect the fast timescale response to laser ablation, they do not affect overall tissue mechanics. In conclusion, our work shows that cellular dynamics and tissue shape changes during *Drosophila* pupal wing morphogenesis are independent of one potential chemical guiding cue, core PCP.

## 1 Introduction

The spatial-temporal pattern of mechanical deformation during tissue morphogenesis is often guided by patterns of chemical signaling. Precisely how chemical signaling couples with the mechanics of morphogenesis, however, remains an active area of research. One known signal that is organized across tissues is the core planar cell polarity (PCP) pathway, a dynamic set of interacting membrane proteins that polarize within the plane of a tissue and globally align to orient structures such as hairs on the fly wing or animal skin (Cetera et al., 2017; Gubb and García-Bellido, 1982; Guo et al., 2004; Strutt and Strutt, 2002) and sterocilia of the vertebrate inner ear (Deans, 2021; Eaton, 1997). In many systems, core PCP also organizes patterns of dynamic cellular behaviors underlying tissue morphogenesis. For example, in zebrafish, non-canonical Wnt-mediated PCP regulates the actomyosin cytoskeleton to coordinate cell rearrangements during heart chamber remodelling (Merks et al., 2018). In *Drosophila*, there is also evidence to suggest that PCP components influence the orientation of cell division and cell shape changes in the larval wing disc, as well as cell rearrangements in the pupal notum (Baena-López et al., 2005; Bosveld et al., 2012; Mao et al., 2011; Ségalen et al., 2010). Here, we examine a potential role for core PCP in the dynamics and mechanics of morphogenesis using the *Drosophila* pupal wing.

The *Drosophila* wing is a flat epithelium that can be imaged at high spatial-temporal resolution *in vivo* during large-scale tissue flows. During the pupal stage the proximal hinge region of the wing contracts and pulls on the blade region, generating mechanical stress that is counteracted by marginal connections mediated by the extracellular protein Dumpy (Etournay et al., 2015; Ray et al., 2015; Wilkin et al., 2000). As a consequence, the tissue elongates along the proximal-distal (PD) axis and narrows along the anterior-posterior (AP) axis to resemble the adult wing (Etournay et al., 2015). Both cell elongation changes and cell rearrangements are important for tissue deformation. To some extent, mechanical stress induces these cell behaviors. However, the reduction of mechanical stress in a *dumpy* mutant does not completely eliminate cell rearrangements, suggesting that there could be other patterning cues that drive oriented cell rearrangements (Etournay et al., 2015). We therefore wondered whether PCP systems could orient cell behaviors, such as cell rearrangements, during pupal wing morphogenesis.

In the *Drosophila* wing, there are two PCP systems termed Fat and core PCP. The Fat PCP system consists of two cadherins Fat and Dachsous, a cytoplasmic kinase Four-jointed and an atypical myosin Dachs (Clark et al., 1995; Ishikawa et al., 2008; Mahoney et al., 1991; Mao et al., 2006). The core PCP system is composed of two transmembrane cadherins Frizzled (Fz) and Flamingo or Starry night (Fmi, Stan), the transmembrane protein Strabismus or Van Gogh (Stbm, Vang) and the cytosolic components Dishevelled (Dsh), Prickle (Pk), and Diego (Dgo) (Adler et al., 1990; Devenport, 2014; Feiguin et al., 2001; Gubb and García-Bellido, 1982; Klingensmith et al., 1994; Taylor et al., 1998; Vinson and Adler, 1987; Wolff and Rubin, 1998). Our group has shown that tissue-scale patterns of PCP emerge during larval stages and then are dynamically reoriented during pupal tissue flows by shear stress (Aigouy et al., 2010; Merkel et al., 2014; Sagner et al., 2012). At the onset of pupal morphogenesis, both systems are margin-oriented, however as morphogenesis proceeds, core PCP reorients to point along the proximal-distal axis, whereas Fat PCP remains margin-oriented until very late, when it reorients towards veins (Fig S1.1A-B) (Merkel et al., 2014). Whether these systems and their reorientation influence tissue mechanics during pupal morphogenesis is unknown.

Here, we examine cellular dynamics in tissues mutant for core PCP and we find that they are largely unperturbed, indicating that core PCP does not have an essential role in organizing global patterns of cell rearrangements in the pupal wing. We also performed an extensive analysis of the mechanics using laser ablation, developing a rheological model to interpret the results. We find that mutants in core PCP differ from wild type in the initial retraction velocity upon laser ablation. We find, however, that this difference is produced from the very fast timescale response, which does not appear to affect morphogenesis and overall tissue stresses, consistent with the lack of phenotype in cellular dynamics.

## 2 Results

### 2.1 Core PCP does not guide cellular dynamics during pupal wing morphogenesis

To investigate the role of core PCP in orienting cell behaviors during pupal wing morphogenesis, we analyzed cell dynamics in wild type (*wt*) and three different *core PCP* mutant tissues: *prickle* (*pk30*), *strabismus* (*stbm6*) and *flamingo* (*fmi*^*frz3*^, aka *stan*^*frz3*^, abbreviated here simply as *fmi*). In *pk30*, the core and Fat PCP systems remain aligned together toward the margin and the magnitude of Stbm polarity is reduced (Merkel et al., 2014). In *stbm6* and *fmi*, the core PCP network is absent (Fig S1.1A-B) (Merkel et al., 2014). We analyzed shape changes of the wing blade during morphogenesis and decomposed these changes into contributions from cell elongation changes and cell rearrangements, which include cell neighbor exchanges, cell divisions, cell extrusions, and correlation effects (Fig 1A-C) (Etournay et al., 2015). Overall, the wing blade elongates along the PD axis by the end of morphogenesis (blue line in Fig 1D). Cells first elongate along the PD axis and then relax to more isotropic shapes (green line in Fig 1D). Cell rearrangements, however, go the opposite direction, initially contributing to AP deformation, before turning around to contribute to PD deformation (magenta line in Fig 1D). We introduce here a relative timescale, where time 0 h is the peak of cell elongation. This new scale allows us to handle variation in the timing of the onset of pupal wing morphogenesis, which we have observed recently (see Appendix 1).

**Figure 1:**
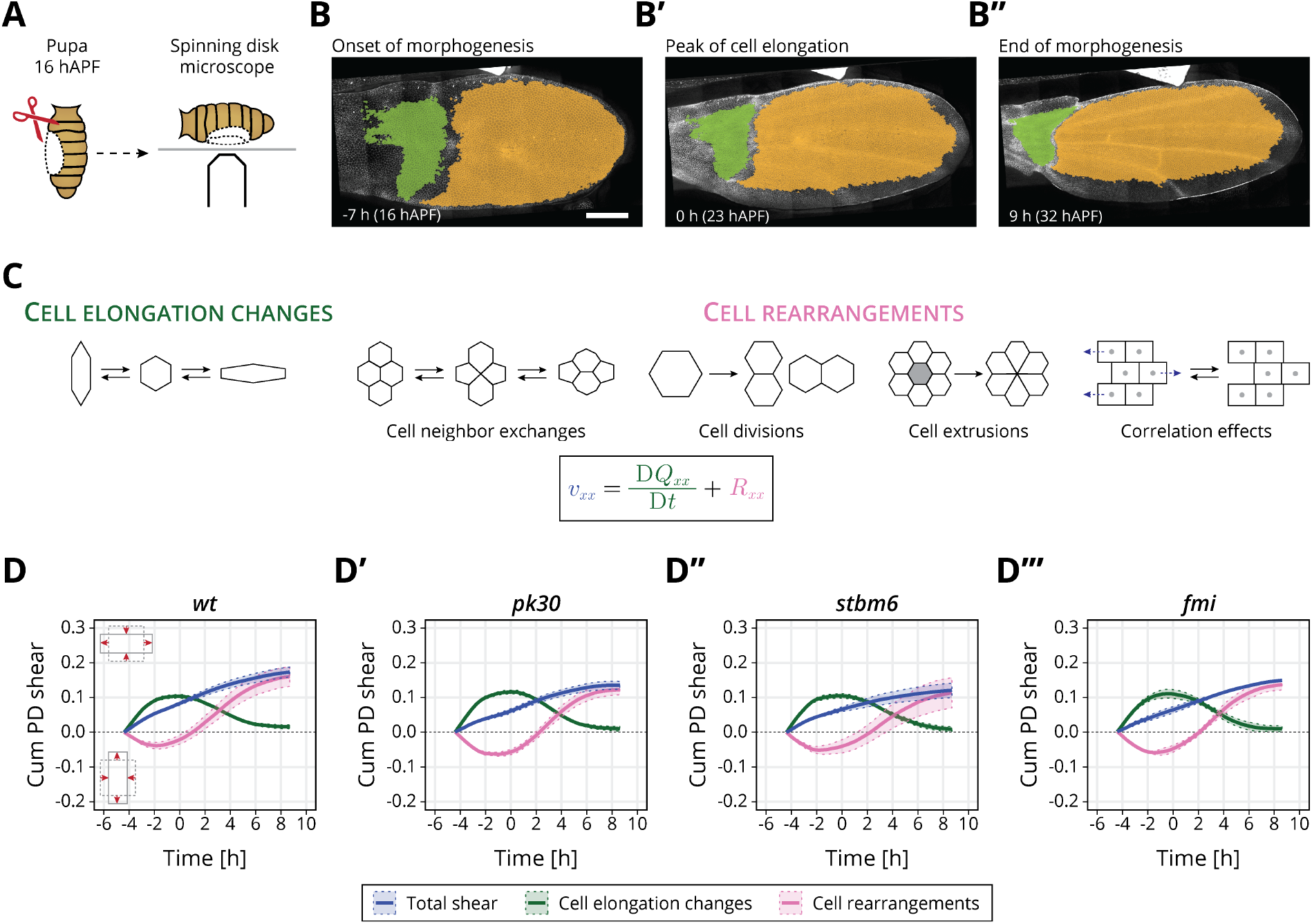
Core PCP does not orient cellular behaviors and tissue reshaping during pupal wing morphogenesis: (A) Cartoon of pupal wing dissection at 16 hAPF and imaging using a spinning disk microscope. (B-B^*′′*^) Images of a *wt* wing at -7, 0 and 9 h (for this movie these times correspond to 16, 23 and 32 hAPF). The green and orange regions correspond to the hinge and blade, respectively. Anterior is up; proximal to the left. Scale bar, 100 *μ*m. (C) Schematic of the cellular contributions underlying anisotropic tissue deformation. Total shear is the sum of cell elongation changes (green) and cell rearrangements (magenta). (D-D^*′′′*^) Accumulated tissue shear during morphogenesis in the blade region averaged for (D) *wt* (n=4), (D^*′*^) *pk30* (n=3), (D^*′′*^) *stbm6* (n=3) and (D^*′′′*^) *fmi* (n=2) movies. Solid line indicates the mean, and the shaded regions enclose *±* SEM. Differences in total accumulated shear are not statistically relevant (Fig S1.2C). The time is relative to the peak of cell elongation. Supplementary data: S4.1.

Interestingly, we find that the overall deformation of the pupal wing blade is unchanged in *core PCP* mutants (Fig 1D-D^*′′′*^). Furthermore, the dynamics of cell elongation changes and the cell rearrangements, when averaged across the entire blade, occur normally. We observe that by the end of the process, only slightly lower total shear has occurred in the *core PCP* mutants, caused by slightly less cell rearrangements, but these subtle changes are not statistically significant (Fig 1D-D^*′′′*^, Fig S1.2C). The cellular dynamics contributing to isotropic tissue deformation also do not show differences between *wt* and *core PCP* mutant tissues (Fig S1.2D).

We also looked for differences in the behavior of regions of the wing blade subdivided along the PD axis (Fig S1.3E-F), as previous work has shown that distal regions of the wing blade shear more at early times, while proximal regions start deforming later (Merkel et al., 2017). Again, we find only subtle differences between *core PCP* mutants and *wt* when we subdivide the wing into regions along the PD axis: proximal blade regions (Region 1, Fig S1.3F) have slightly more AP-oriented cell rearrangements and end up with less overall total shear, whereas distal regions (Region 4, Fig S1.3F) shear slightly faster due to more PD-oriented cell elongation. These differences, however, are not statistically significant (Fig S1.4G). From this analysis, we conclude that core PCP does not guide the global patterns of cell dynamics during pupal morphogenesis. The subtle reduction in shear observed in the mutants is nevertheless consistent with the slight defect in adult wing shape: *pk30* and *stbm6* (but not *fmi*) mutant wings are slightly rounder and wider than *wt* (Fig S1.5H). These differences between *core PCP* mutants and *wt* may be detectable in adult wings because we can analyze many more wings, whereas the acquisition of timelapses is technically challenging and time-consuming, and we could only reasonably analyze 2-4 wings per genotype. Alternatively, core PCP may affect later stages of wing development, after the after the pupal stages observed here. To investigate whether the slight differences we observe in the pupal shear patterns and adult wing shape could indicate a more subtle effect of core PCP on cell and tissue mechanics, we next analyzed mechanical stress and strain in a small region of the wing blade.

### 2.2 A rheological model for the response to laser ablation

We investigated cell and tissue mechanics in *core PCP* mutants using laser ablation in a small region of the wing blade. We used a region located between the second and third sensory organs in the intervein region between the L3 and L4 longitudinal veins, which is a region that is easy to identify throughout morphogenesis (Fig 2A). We cut 3-4 cells in a line along the AP axis and measure the displacement of the tissue (Fig 2A, Supplemental movie 1). Previously, we reported that initial recoil velocity measured along the PD axis in *wt* peaks around -8 h (20 hAPF in Iyer et al., 2019), and therefore we first focus on this timepoint. We find that *core PCP* mutants have significantly lower recoil velocity (Fig 2B, Fig S2.1A), suggesting that there is a mechanical defect in these mutants.

**Figure 2:**
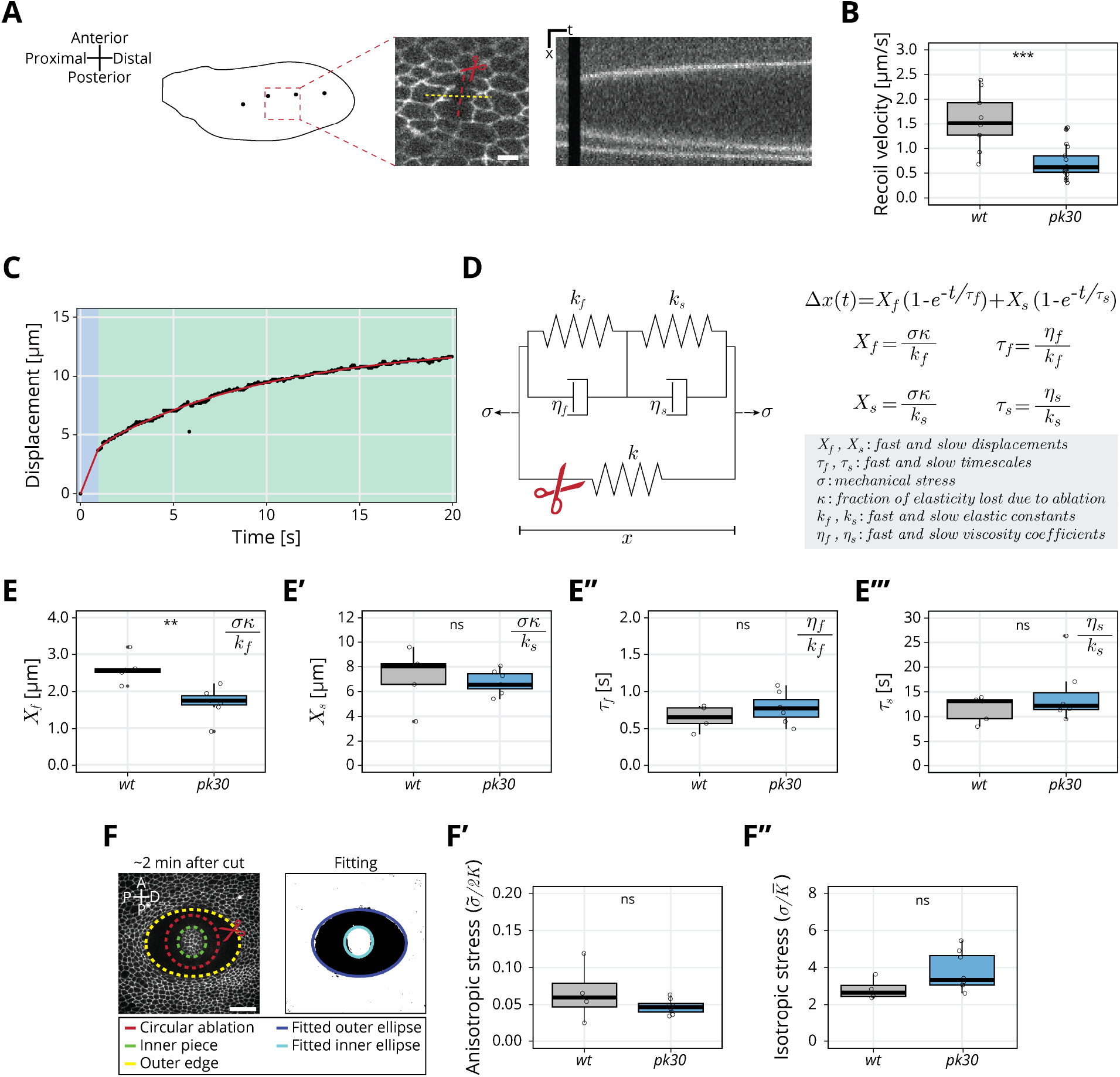
Rheological model for the response to laser ablation: (A) Schematic of a *wt* wing at -8 h. Linear laser ablation experiments were performed in the blade region enclosed by the red square. Dots on the wing cartoon indicate sensory organs. The red line corresponds to the ablation and the kymograph was drawn perpendicularly to the cut (yellow). Scale bar, 5 *μ*m. (B) Initial recoil velocity upon ablation along the PD axis at -8 h for *wt* (gray) and *pk30* (blue) tissues (n⩾9). Significance is estimated using the Mann–Whitney U test. ***, p-val⩽0.001. (C) Example of the measured displacement after laser ablation (black dots) and corresponding exponential fit of the mechanical model (red curve). The blue and green regions highlight the displacement in the fast and slow timescale, respectively. (D) Description of the mechanical model that was devised to analyze the tissue response upon laser ablation. After the cut, the spring with elastic constant *k* is ablated (red scissor), and the tissue response is given by the combination of the two Kelvin-Voigt models arranged in series. These two correspond to the fast response given by *k*_*f*_ and *η*_*f*_ and the slow response given by *k*_*s*_ and *η*_*s*_. The mechanical stress *σ*is constant. The membrane displacement Δ*x*(*t*) is calculated as a sum of the displacement (*X*_*f*_) associated with the fast timescale (*τ*_*f*_) and the displacement (*X*_*s*_) associated with the slow timescale (*τ*_*s*_). (E-E^*′′′*^) Values obtained for each of the four fitting parameters when fit to the data. (E) Displacement associated with the fast and (E^*′*^) slow timescale for *wt* (gray) and *pk30* (blue). (E^*′′*^) Fast and (E^*′′′*^) slow timescale for *wt* (gray) and *pk30* (blue) (n⩾5). Significance is estimated using the Student’s t-test. **, p-val⩽0.01; ns, p-val*>*0.05. (F) Example of a circular laser ablation used for analysis with ESCA. The left image shows the final shape of the ablation around 2 min after cut, and the right image shows the corresponding segmented image, where the inner and outer pieces were fit with ellipses. After the fitting, the model outputs the anisotropic and isotropic stress (equations shown on the right side). Scale bar, 20 *μ*m. A=Anterior, P*=posterior, D=distal, P=proximal. (F^*′*^) Anisotropic stress 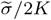 for *wt* (gray) and *pk30* (blue) tissues at -8 h (n⩾4). Significance is estimated using the Mann–Whitney U test. ns, p-val*>*0.05. (F^*′′*^) Isotropic stress 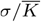 for *wt* (gray) and *pk30* (blue) tissues at -8 h (n⩾4). Significance is estimated using the Mann–Whitney U test. ns, p-val*>*0.05. The time is relative to the peak of cell elongation. In all plots, each empty circle indicates one cut, and the box plots summarize the data: thick black line indicates the median; the boxes enclose the 1st and 3rd quartiles; lines extend to the minimum and maximum without outliers, and filled circles mark outliers. Supplementary data: S4.2.

As initial recoil velocity is often used as a proxy for mechanical stress (Etournay et al., 2015; Farhadifar et al., 2007; Iyer et al., 2019; Mayer et al., 2010), this result seems to suggest that the PCP mutant wings generate less mechanical stress during morphogenesis, even though the cellular dynamics is basically unperturbed. To explore this phenotype in more detail, we considered that the response to laser ablation is not exactly a direct measure of mechanical stress, as it is also affected by cellular material properties. We thus further analyzed the full kinetics of the linear laser ablations, focusing on the *pk30* mutant, and developed a rheological model to interpret the results. When plotting displacement of the nearest bond to ablation over time, we observe both a fast (<1 s) and a slow (<20 s) regime (Fig 2C). We therefore developed a model consisting of two Kelvin-Voigt (KV) elements in series (Fig 2D) to represent the tissue after ablation. The two KV elements have different elastic constants (*k*_*f*_ and *k*_*s*_) and viscosities (*η*_*f*_ and *η*_*s*_). Before ablation, the system is subjected to a constant stress (*σ*) and contains a spring with elastic constant *k*, which represents the cell patch that will be ablated. Upon ablation, the third spring is removed which leads to change in strain of our rheological model. We represent this strain by a displacement Δ*x* as a function of time given by

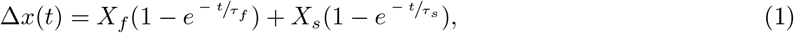

where *X*_*f*_ = *σκ/k*_*f*_ is the displacement associated with the fast timescale, *τ*_*f*_ = *η*_*f*_ */k*_*f*_, and *X*_*s*_ = *σκ/k*_*s*_ is the displacement associated with the slow timescale, *τ*_*s*_ = *η*_*s*_*/k*_*s*_. Here, 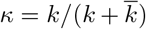 is the fraction of the overall system elasticity lost due to ablation (see Materials and Methods 5.6.3) and 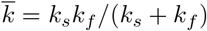 is the elasticity of the two KV elements connected in series. With this model, we presume the properties of the ablated cell itself, including its membrane, adhesion proteins, and actomyosin cortex likely dominate the fast timescale response. The slow timescale response is a collective effect emerging from the ablated cell together with its surrounding cellular network.

We analyzed experimentally measured displacement over time for each ablation and then fit the data to our model with four parameters (*X*_*f*_, *X*_*s*_, *τ*_*f*_ and *τ*_*s*_) (Eq 1, Fig 2E-E^*′′′*^). Surprisingly, we find that the only parameter that changes between *pk30* and *wt* is *X*_*f*_, the displacement associated with the fast timescale (Fig 2E-E^*′′′*^).

To interpret this result, we consider that these four fitted parameters constrain the five mechanical model parameters (Fig 2D) but do not provide a unique solution. Since only one measured parameter changes, we asked what is the simplest set of model parameter changes that could have such an effect. To this end, we first note that the measured values of *X*_*f*_ and *X*_*s*_ (1.8 − 2.6 *μm* vs 6 − 8 *μm*, respectively) indicate *k*_*f*_ ≫ *k*_*s*_ and therefore the overall elasticity of our rheological model is largely determined by the elasticity of the slow relaxation 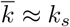. If we also consider that the contribution to the elasticity of the cellular patch from the ablated cells, represented by *k* in the model, is small, then we can approximate *κ* ≈ *k/k*_*s*_ and therefore *X*_*f*_ ≈ *σk/*(*k*_*s*_*k*_*f*_), (see Materials and Methods 5.6.2). To probe whether a change of *σ/k*_*s*_ can account for the change of *X*_*f*_ in the *pk30* mutant, we used ESCA (Elliptical Shape after Circular Ablation) (Dye et al., 2021) to infer anisotropic and isotropic stress in the tissue from the final relaxed tissue shape after a circular ablation (Fig 2F, see Materials and Methods 5.6.2). We find no significant difference between *wt* and *pk30* mutants (Fig 2F-F^*′′*^), therefore we conclude that *σ/k*_*s*_ cannot account for the observed change in *X*_*f*_. Furthermore, we find from ESCA that the ratio of shear and isotropic elastic constants also does not change between *wt* and *pk30* mutant, suggesting that *σ* does not change (Fig S2.1B).

Therefore, we can account for the observed changes in the *pk30* mutant with a change of only one elastic coefficient *k*_*f*_. This result suggests that a change in *k*_*f*_ is inversely proportional to the observed change in *X*_*f*_. In this scenario, the only other parameter that changes is *η*_*f*_ such that *τ*_*f*_ does not change, as observed. The conclusion that only the short time response to the ablation is affected in the *pk30* mutant is consistent with the lack of a clear phenotype in the large-scale tissue flows.

### 2.3 Dynamics of stress and cell elongation throughout morphogenesis in wild type and *pk30* mutant

To examine the effect of PCP mutation throughout pupal morphogenesis, we aimed to simplify the time intensive segmentation of the full ablation dynamics. To this end, we measured the initial recoil velocity *v* defined by the displacement after ablation Δ*x* over a short time interval *δt* = 0.65 *s*. The recoil velocity can be expressed in terms of our model parameters as 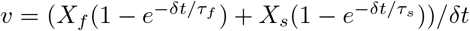. Since the value of *δt* is comparable to the fast time-scale *τ*_*f*_, about 63% of the *X*_*f*_ value relaxes over that time, while at the same time only about 5% of the *X*_*s*_ value is relaxed. Using the measured values of *X*_*f*_ and *X*_*s*_, we estimate that the fast timescale dynamics contributes about 80% of the *v* value. Therefore, the initial retraction velocity is a good proxy for the fast displacement *X*_*f*_.

We find that the initial recoil velocity along the PD axis peaks at -8 h before declining again by 4 h (Fig 3A), consistent with previous work (Iyer et al., 2019). The behaviour of the recoil velocity in the *pk30* mutant is qualitatively similar throughout morphogenesis, however, with significantly lower magnitude than *wt* (Fig 3A). We also observed this behavior in *stbm6* and *fmi* mutant tissues (Fig S3.1A). This result indicates that *X*_*f*_ is lower in *core PCP* mutants than in *wt* throughout morphogenesis. We also performed ESCA over time in *pk30* mutants and observe that anisotropic stress 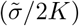 rises early during morphogenesis before eventually declining (Fig 3B), whereas isotropic stress 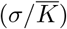 remains fairly constant (Fig 3B^*′*^). Importantly, ESCA does not report any difference in measured stresses between *pk30* and *wt*, nor in the ratio of elastic constants (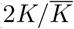, Fig S3.2), indicating that our conclusions based on the -8 h timepoint are true throughout morphogenesis, and differences in *σ/k*_*s*_ between *pk30* and *wt* do not account for differences in *X*_*f*_.

**Figure 3:**
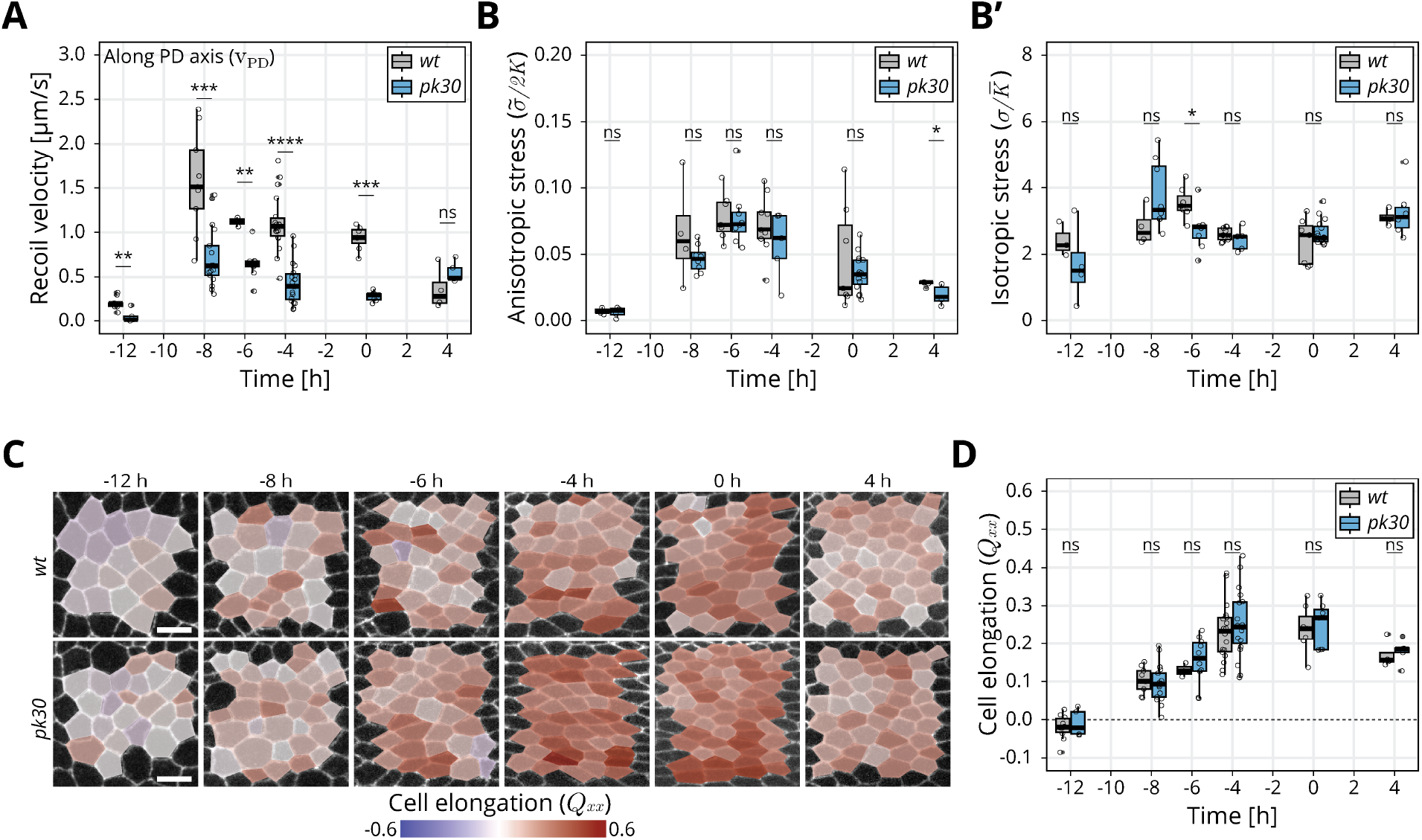
Dynamics of stress and cell elongation throughout morphogenesis in *wt* and *pk30* mutant: (A) Initial recoil velocity upon ablation along the PD axis throughout morphogenesis for *wt* (gray) and *pk30* (blue) tissues (n⩾3). Significance is estimated using the Mann–Whitney U test. ****, p-val⩽0.0001; ***, p-val⩽0.001; **, p-val⩽0.01; ns, p-val*>*0.05. (B) ESCA results for anisotropic stress 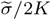 for *wt* (gray) and *pk30* (blue) tissues throughout morphogenesis (n⩾3). Significance is estimated using the Mann–Whitney U test. *, p-val*<*0.05; ns, p-val*>*0.05. (B^*′*^) ESCA results for isotropic stress 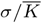 for *wt* (gray) and *pk30* (blue) throughout morphogenesis (n⩾3). Significance is estimated using the Mann–Whitney U test. *, p-val*<*0.05; ns, p-val*>*0.05. (C) Color-coded PD component of cell elongation *Q*_*xx*_ in the blade region between the second and third sensory organs found in the intervein region between L2 and L3. The images correspond to *wt* (top row) and *pk30* (bottom row) wings throughout morphogenesis. Scale bar, 5 *μ*m. (D) Quantification of *Q*_*xx*_ in this region throughout morphogenesis for *wt* (gray) and *pk30* (blue) (n⩾3). Significance is estimated using the Mann–Whitney U test. ns, p-val*>*0.05. The time is relative to the peak of cell elongation. In all plots, each empty circle indicates one experiment, and the box plots summarize the data: thick black line indicates the median; the boxes enclose the 1st and 3rd quartiles; lines extend to the minimum and maximum without outliers, and filled circles mark outliers. Supplementary data: S4.3.

To directly compare linear laser ablations to ESCA, we performed linear ablations also in the perpendicular orientation. We then compared the difference in retraction velocities in these two orientations with the anisotropic tissue stress measured by ESCA (Fig S3.1B-C). We find that the magnitude of the response is lower in *pk30* compared to *wt*, as observed for the PD orientation alone. Interestingly, the dynamics of the response in *wt* is comparable to the dynamics of anisotropic stress measured by ESCA (Fig 3B, Fig S3.1C). However, for *pk30*, the dynamics reflected by initial recoil velocity is qualitatively different from the stress measured by ESCA (Fig 3B, Fig S3.1C), indicating that *k*_*f*_ could be changing in time in *pk30* mutants.

To further probe the possible role of Prickle in epithelial mechanics, we also measured the dynamics of the proximal-distal component of cell elongation (*Q*_*xx*_) in *wt* and *pk30* (Fig 3C). Interestingly, in both cases, anisotropic stress peaks around -6 h (Fig 3B), whereas *Q*_*xx*_ peaks significantly later, between -4 h and 0 h, indicating that active cellular stresses contributing to cell elongation change in time. However, there are no differences between *wt* and *pk30* (Fig 3D), showing that core PCP also does not affect active stresses underlying the dynamics of cell elongation during pupal morphogenesis.

## 3 Discussion

Here, we used the *Drosophila* pupal wing as a model for studying the interplay between planar polarized chemical signaling components, specifically the core PCP pathway, and the mechanical forces underlying tissue morphogenesis. Unlike in other systems, where core PCP seems to organize patterns of cellular behaviors (Bosveld et al., 2012; Ségalen et al., 2010; Shindo and Wallingford, 2014, reviewed in Wang and Nathans, 2007), here an extensive analysis of core PCP mutants shows no significant phenotype in pupal wing morphogenesis. We find no differences in overall tissue shape change, nor in the pattern or dynamics of underlying cellular contributions. Furthermore, we found no significant differences in tissue mechanical stress or in cell elongation over time. Generally these results are consistent in mutants that destroy core PCP polarity (*stbm6* and *fmi*) or prevent its decoupling from Fat (*pk30*).

Interestingly, we do observe a phenotype in the initial retraction velocity upon laser ablation between *pk30* and *wt*. Retraction velocity after a laser ablation is often used a proxy for tissue mechanical stresses. If the differences between the *wt* and *core PCP* mutant wings observed in the retraction velocity indeed reflect the difference in mechanical stress, how can we understand lack of phenotype in tissue shape changes, cell dynamics and cell elongation? A more detailed analysis suggests that the difference between the two genotypes arises from a difference in the elastic constant *k*_*f*_ underlying the fast timescale response (*τ*_*f*_ = 0.65 *s*) to the ablation. Our results are consistent with the initial retraction velocity being proportional to tissue stress, however, the proportionality factor can depend on the genotype and can change in time (Fig S3.1C). This result highlights a limitation of comparing tissue mechanics in different genotypes based on retraction velocity measurements.

What is the the biophysical nature of the fast response to laser ablation? We hypothesize that processes that react on time-scales *<* 1 *s* could be related to cortical mechanics of cell bonds or possibly changes in cell hydraulics after an ablation. In our simple model, these processes affect the stiff spring *k*_*f*_, which has a small contribution to the effective cell and tissue elasticity 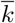, which is dominated by the soft spring *k*_*s*_, see Results 2.2. This could explain why the differences we observed at the fast time scales in *wt* and *pk30* wings are not reflected in wing morphogenesis. However, it remains unclear how core PCP affects only the elasticity *k*_*f*_ of the fast response and it remains an open possibility that there is a mechanism that compensates for such a change so that there is no change at larger scales.

While we have shown that core PCP is not required to organize the dynamic patterns of cellular events underlying pupal wing morphogenesis, it might still affect later stages of wing development. Furthermore, there may still be other patterning systems acting redundantly or independently with core PCP. For example, the Fat PCP system and Toll-like receptors have been shown to influence the orientation of cellular rearrangements and cell divisions in other contexts (Lavalou et al., 2021; Mao et al., 2006; Paré et al., 2014, reviewed in Umetsu, 2022). Whether and how other polarity systems influence pupal wing morphogenesis remains unknown.

## Acknowledgements

We thank Stephan Grill for giving us access to the microscope used for laser ablation. We thank the Light Microscopy Facility, the Computer Department, and the Fly Keepers of the MPI-CBG for their support and expertise. We would like to thank Christian Dahmann and Jana Fuhrmann for comments on the manuscript prior to publication. This work was funded by Germany’s Excellence Strategy – EXC-2068 – 390729961– Cluster of Excellence Physics of Life of TU Dresden, as well as grants awarded to SE from the Deutsche Forschungsgemeinschaft (SPP1782, EA4/10-1, EA4/10-2) and core funding of the Max-Planck Society to SE. NAD additionally acknowledges funding from the Deutsche Krebshilfe (MSNZ P2 Dresden). AK and RPG were funded through the Elbe PhD program. FSG was supported by a DOC Fellowship of the Austrian Academy of Sciences. CD acknowledges the support of a postdoctoral fellowship from the LabEx “Who Am I?” (ANR-11-LABX-0071) and the Université Paris Cité IdEx (ANR-18-IDEX-0001) funded by the French Government through its “Investments for the Future”. We dedicate this work to our coauthor Prof. Dr. Suzanne Eaton, who tragically passed away before the finalization of the project.

## Author Contributions

**RPP**: Investigation, Formal analysis, Software, Validation, Data curation, Writing - original draft, Writing - Review and editing, Visualization. **FSG**: Investigation, Formal analysis, Software, Validation, Data curation, Writing - Review and editing. **AK**: Methodology, Formal analysis, Software, Writing - Review and editing. **CD**: Methodology, Writing - Review and editing **CDM**: Methodology, Resources, Supervision, Writing - Review and editing. **MP**: Methodology, Supervision, Writing - original draft, Writing - Review and editing. **FJ**: Conceptualization, Methodology, Resources, Supervision, Writing - Review and editing, Funding acquisition, Project administration. **NAD**: Resources, Supervision, Writing - original draft, Writing - Review and editing, Data curation, Funding acquisition, Project administration. **SE**: Conceptualization, Methodology, Resources, Supervision, Funding acquisition, Project administration.

## 4 Supplementary Data

### 4.1 Fig 1 Supplementary

**Figure S1.1:**
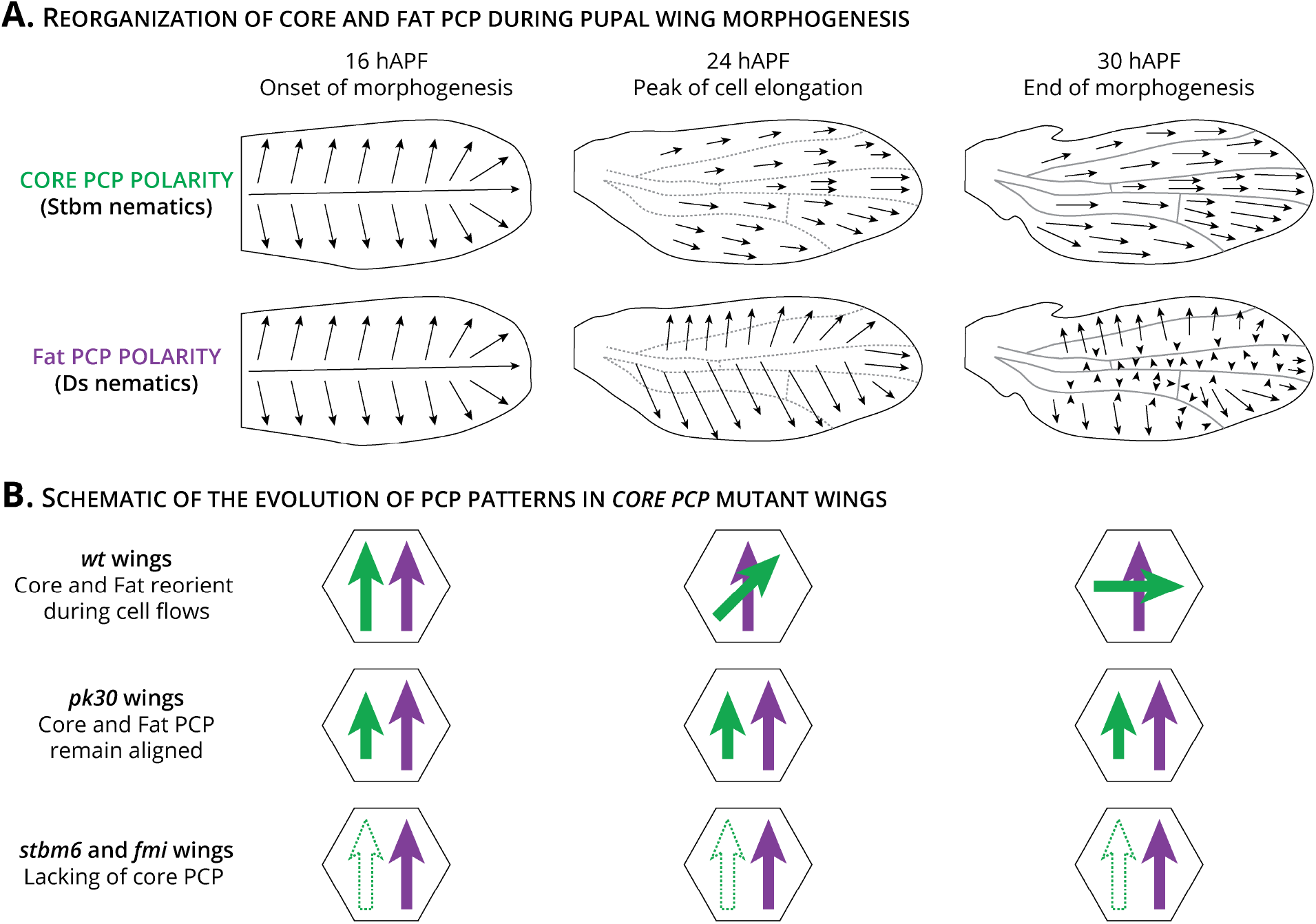
Reorganization of the core and Fat PCP systems during pupal wing morphogenesis: (A) Evolution of the PCP network during pupal wing morphogenesis. Core PCP polarity is based on Stbm::YFP nematics. Initially, core PCP polarity is radially organized towards the wing margin. As tissue flows occur, it reorients towards the distal tip. Fat PCP polarity is based on the pattern of Ds::EGFP. Fat PCP is initially also radially organized. Tissue flows reorganize Fat PCP and by the end of morphogenesis it is perpendicularly oriented to core PCP. Cartoon adapted from Merkel et al. (2014). (B) Schematic of the core (green arrow) and Fat (purple arrow) PCP patterns in *wt, pk30*, and *stbm6* and *fmi* wings. During pupal tissue flows in *wt* wings, core PCP reorients towards the distal tip of the wing. By the end of morphogenesis, core and Fat PCP are perpendicularly aligned. In *pk30* mutant wings, core and Fat PCP remain aligned and core polarity is reduced. In *stbm6* and *fmi* wings, the core PCP network is absent (empty green arrow), while the Fat PCP pattern is unperturbed (purple arrow) (Merkel et al., 2014).

**Figure S1.2:**
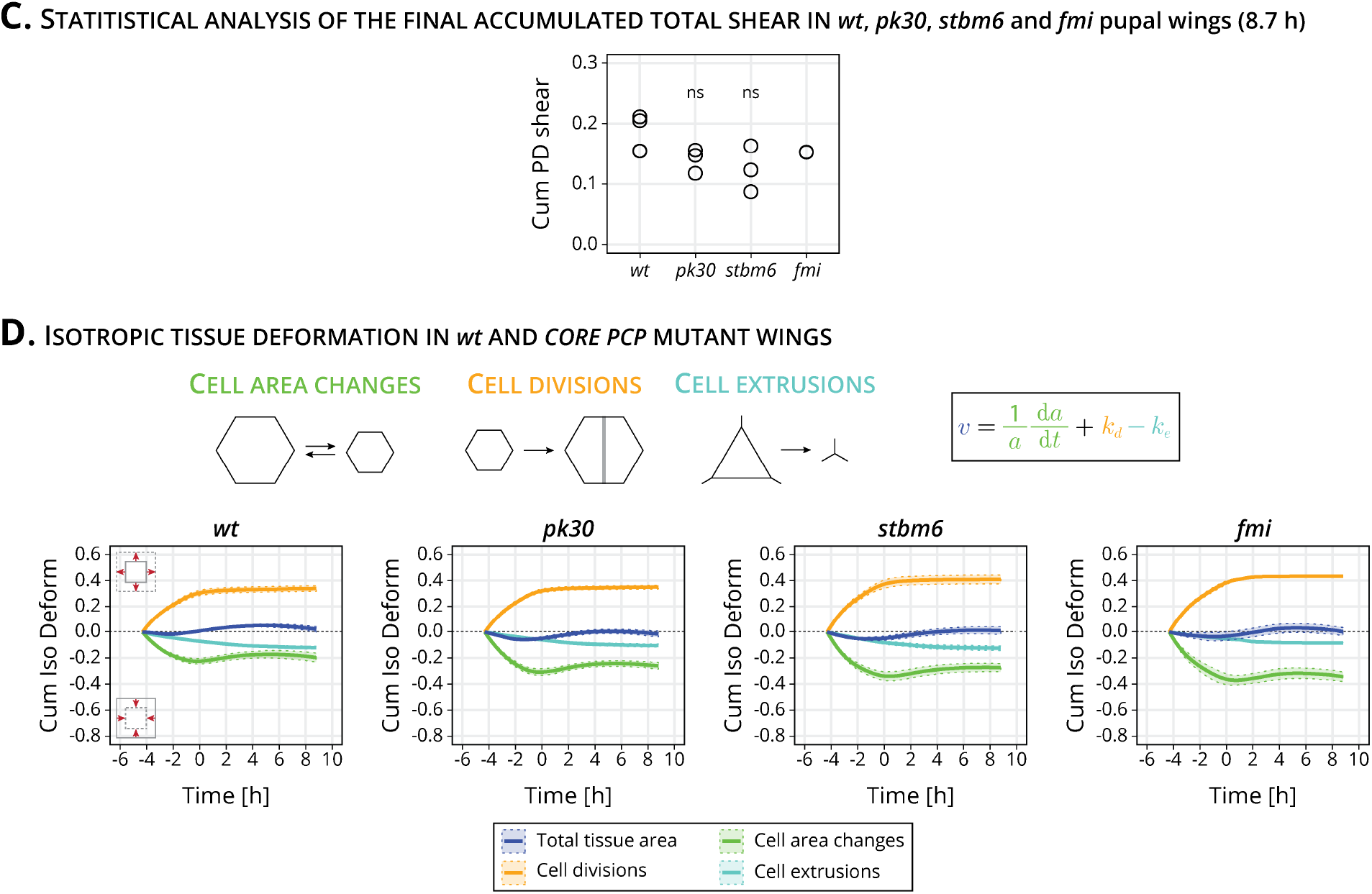
Quantification of final pupal tissue deformation and cellular contributions to isotropic tissue area: (C) Statistical analysis of the final pupal accumulated total tissue shear in *wt* (n=4), *pk30* (n=3), *stbm6* (n=3) and *fmi* (n=2) movies. Significance is estimated using the Kruskal–Wallis test. ns, p-val>0.05. (D) Quantification of isotropic tissue deformation in *wt* (n=4), *pk30* (n=3), *stbm6* (n=3) and *fmi* (n=2) movies. The cellular contributions are cell area changes (green), cell divisions (yellow) and cell extrusions (cyan). Solid line indicates the mean, and the shaded regions enclose *±* SEM. The time is relative to the peak of cell elongation.

**Figure S1.3:**
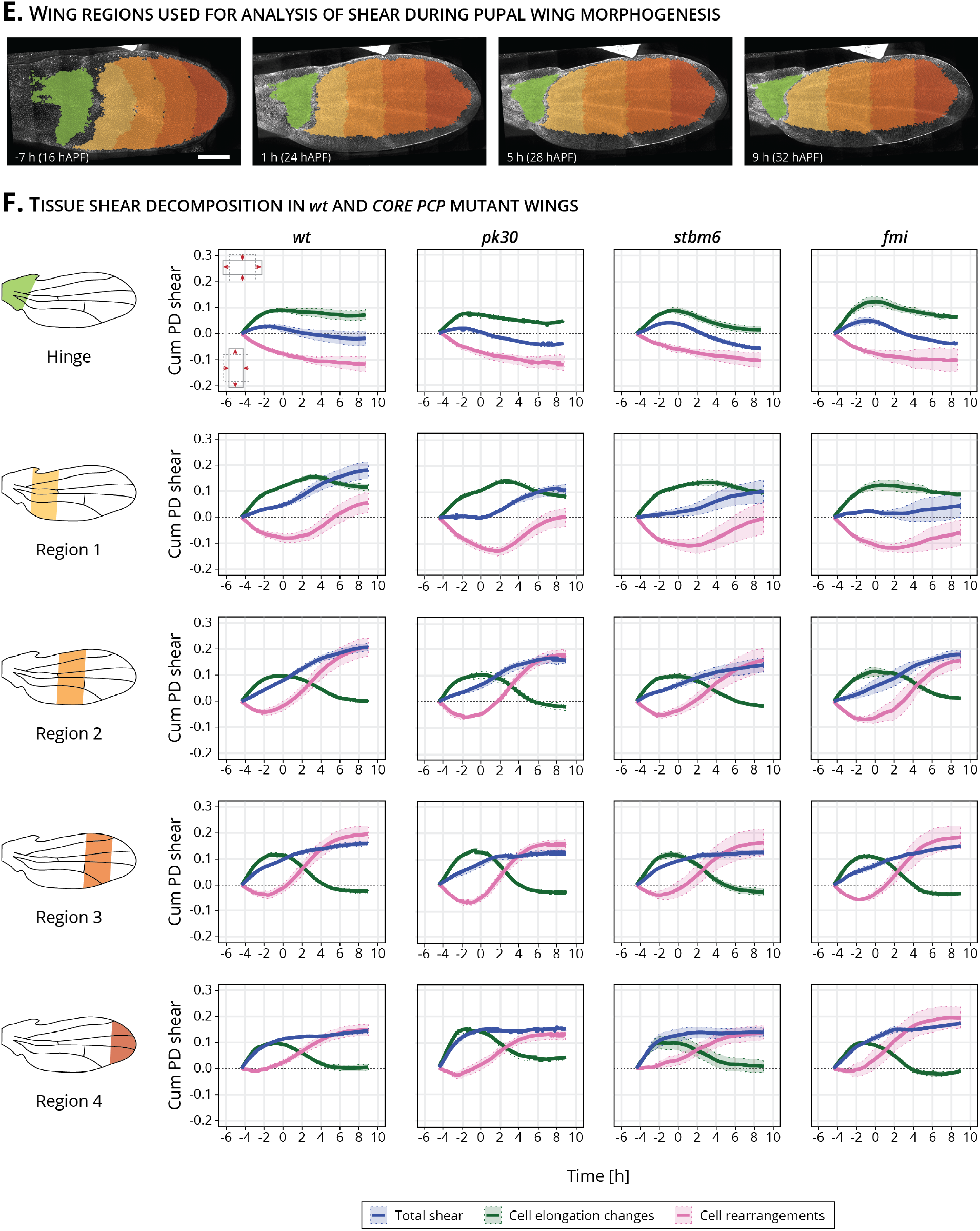
Subregional analysis of tissue shear in the hinge and four blade subregions: (E) Images of a *wt* wing at -7, 1, 5 and 9 h (for this movie it corresponded to 16, 24, 28 and 32 hAPF). The green region corresponds to the hinge and the four blade subregions are shown in an orange color palette. Scale bar, 100 *μ*m. (F) Total shear (dark blue curve) and decomposition into cell elongation changes (green curve) and cell rearrangements (magenta curve) for the hinge and four blade subregions for *wt* (n=4), *pk30* (n=3), *stbm6* (n=3) and *fmi* (n=2). Solid line indicates the mean, and the shaded regions enclose *±* SEM. The time is relative to the peak of cell elongation.

**Figure S1.4:**
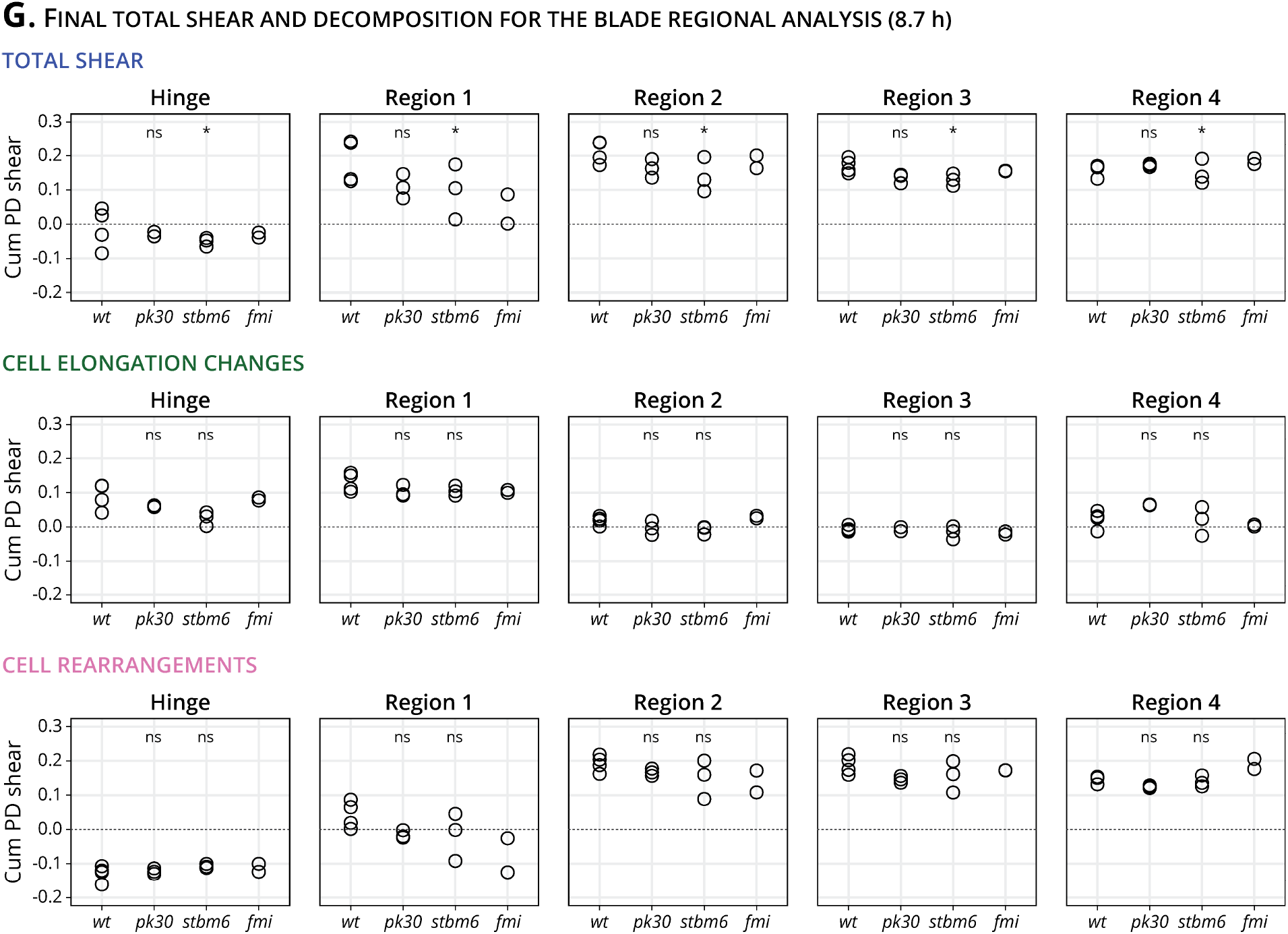
Statistics of final shear in the blade subregions and its cellular contributions.: (G) Quantification of the final total shear (top row), shear caused by cell elongation changes (middle row) and cell rearrangements (bottom row) in the hinge (left column) and four blade subregions for *wt* (n=4), *pk30* (n=3), *stbm6* (n=3) and *fmi* (n=2). Significance is estimated using the Kruskal–Wallis test. *, p-val⩽0.05; ns, p-val*>*0.05.

**Figure S1.5:**
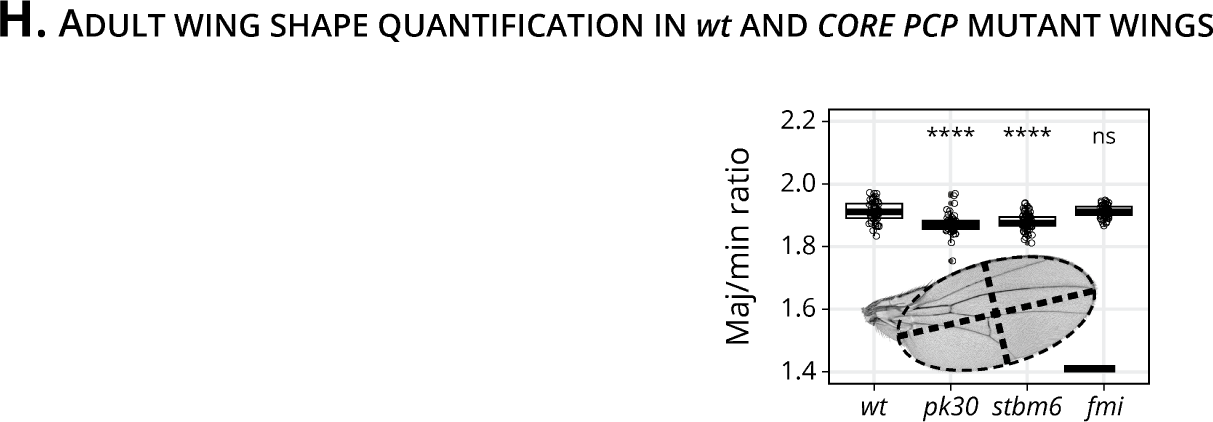
Adult wing shape quantification: (H) Quantification of the adult wing blade major (maj) to minor (min) ratio for *wt, pk30, stbm6* and *fmi* wings (n⩾47). Scale bar, 500 *μ*m. Each empty circle indicates one wing, and the box plots summarize the data: thick black line indicates the median; the boxes enclose the 1st and 3rd quartiles; lines extend to the minimum and maximum without outliers, and filled circles mark outliers. Significance is estimated using the Kruskal–Wallis test. ****, p-val⩽0.0001; ns, p-val*>*0.05.

### 4.2 Fig 2 Supplementary Figures

**Figure S2.1:**
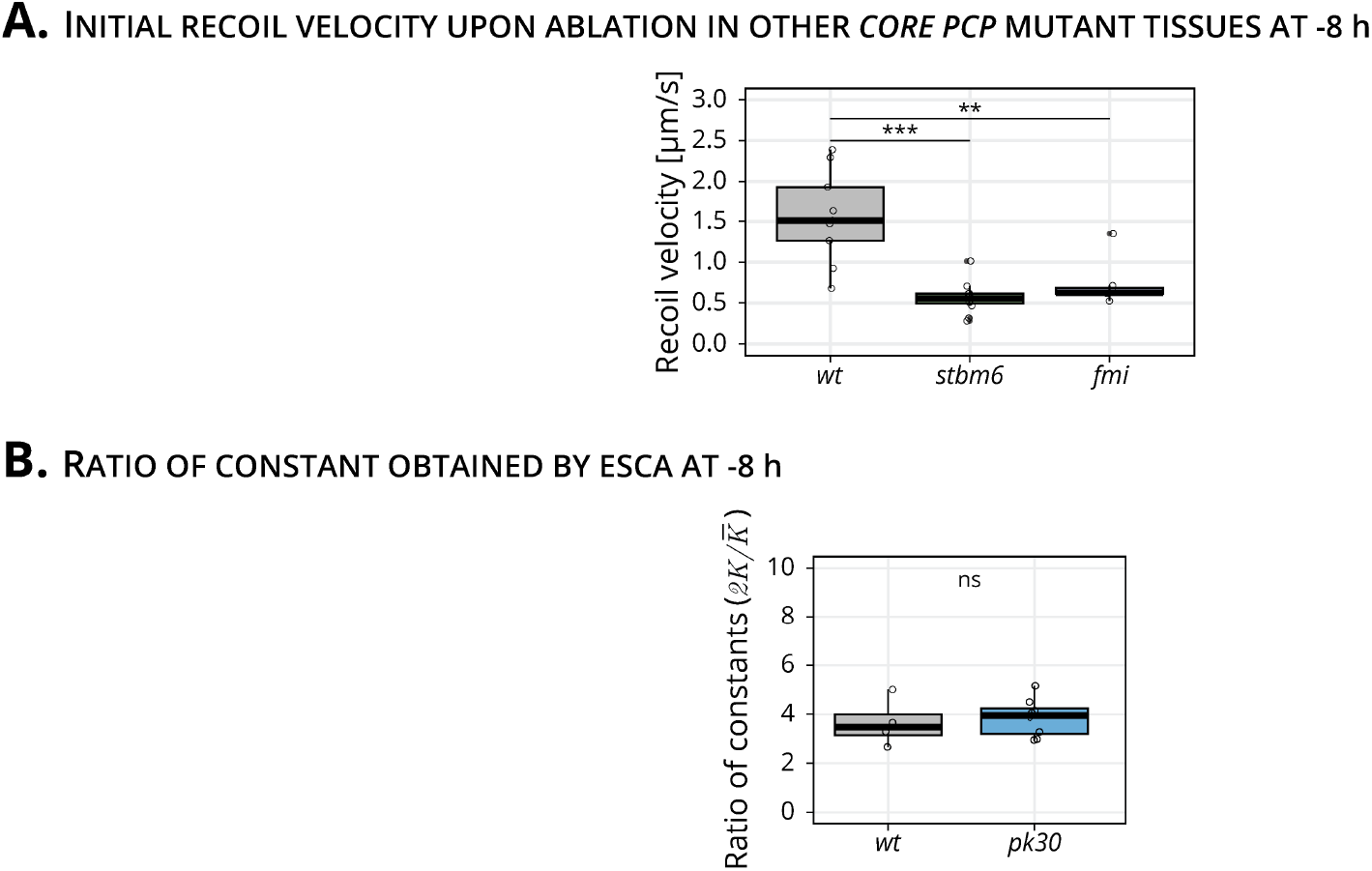
Initial recoil velocity upon linear laser ablation for *stbm6* and *fmi* mutant wings and ratio of elastic constants obtained by ESCA at -8 h: (A) Initial recoil velocity upon ablation along the PD axis for *wt* (gray), *stbm6* (green) and *fmi* (purple) mutant wings at -8 h (n⩾9). Significance is estimated using the Kruskal-Wallis test. ***, p-val⩽0.001; **, p-val⩽0.01. (B) Ratio of elastic constants 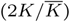 for *wt* and *pk30* (blue) at -8 h (n⩾4). Significance is estimated using the Mann–Whitney U test. ns, p-val*>*0.05. The time is relative to the peak of cell elongation. In both plots, each empty circle indicates one cut, and the box plots summarize the data: thick black line indicates the median; the boxes enclose the 1st and 3rd quartiles; lines extend to the minimum and maximum without outliers, and filled circles mark outliers.

### 4.3 Fig 3 Supplementary Figures

**Figure S3.1:**
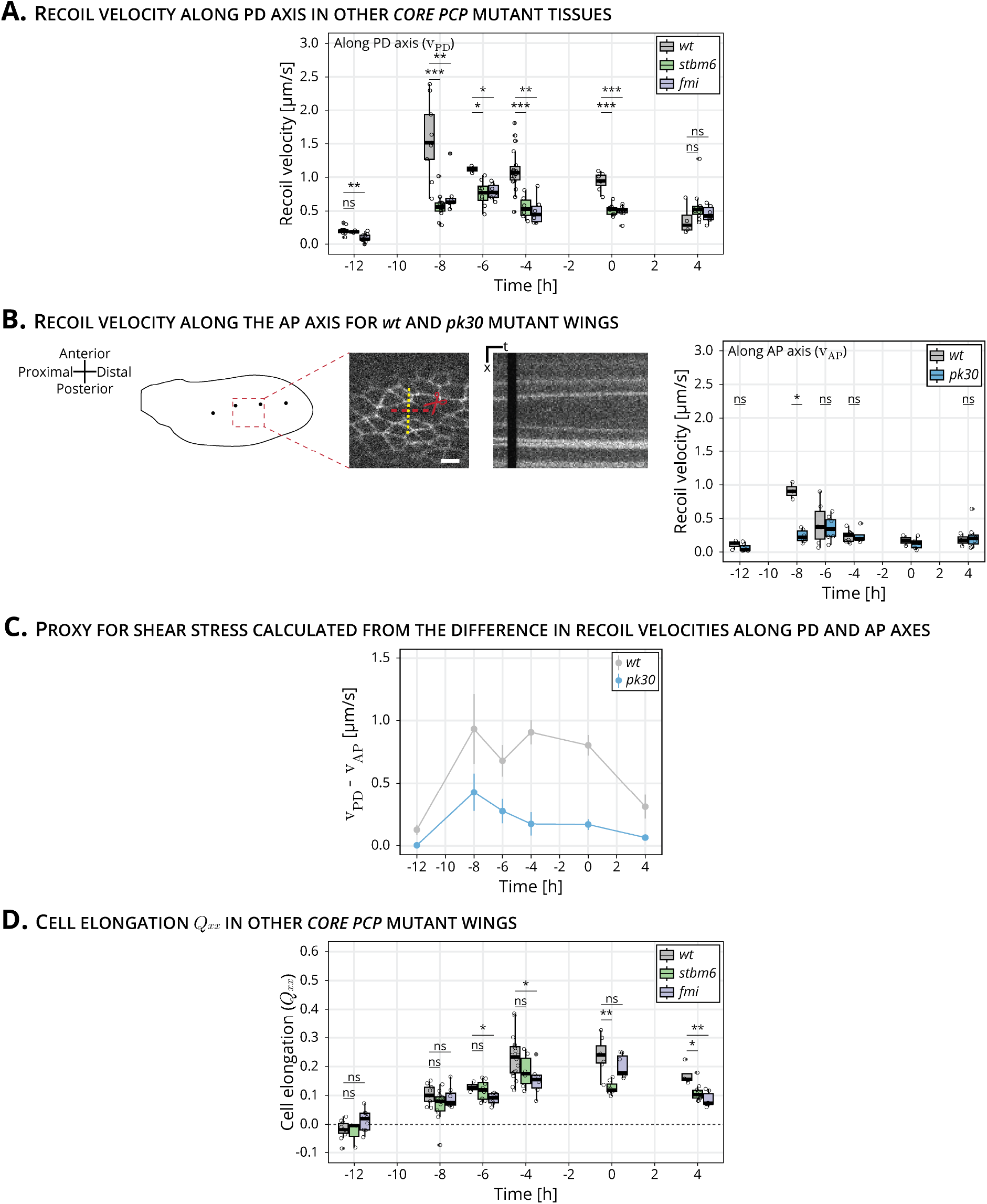
Study of pupal wing mechanics over time: (A) Initial recoil velocity upon ablation along the PD axis for *wt* (gray), *stbm6* (green) and *fmi* (purple) mutant wings throughout morphogenesis (n⩾4). Significance is estimated using the Kruskal-Wallis test. ****, p-val⩽0.0001; ***, p-val⩽0.001; **, p-val⩽0.01; *, p-val⩽0.05; ns, p-val*>*0.05. (B) Left: Schematic of a *wt* wing at -8 h. Linear laser ablation experiments were performed in the blade region enclosed by the red square. Dots on the cartoon indicate sensory organs. Red line corresponds to the ablation; the kymograph was drawn perpendicularly to the cut (yellow). Scale bar, 5 *μ*m. Right: Initial recoil velocity upon ablation along the AP axis for *wt* (gray) and *pk30* (blue) mutant wings throughout morphogenesis (n⩾3, n=2 in *wt* wings at 0 h). Significance is estimated using the Mann–Whitney U test. *, p-val⩽0.05; ns, p-val*>*0.05. (C) Proxy for shear stress calculated as the difference between the initial recoil velocity along the PD (v_PD_) and AP (v_AP_) axes for *wt* (gray) and *pk30* (blue) mutant wings. Filled colored dots correspond to the mean value, and the error bars report the SEM. (D) Quantification of *Q*_*xx*_ in the blade subregion throughout morphogenesis for *wt* (gray), *stbm6* (green) and *fmi* (purple) mutant wings (n⩾4). Significance is estimated using the Kruskal-Wallis test. **, p-val⩽0.01; *, p-val⩽0.05; ns, p-val*>*0.05. The time is relative to the peak of cell elongation. In (A), (B) and (D), each empty circle indicates one experiment, and the box plots summarize the data: thick black line indicates the median; the boxes enclose the 1st and 3rd quartiles; lines extend to the minimum and maximum without outliers, and filled circles mark outliers.

**Figure S3.2:**
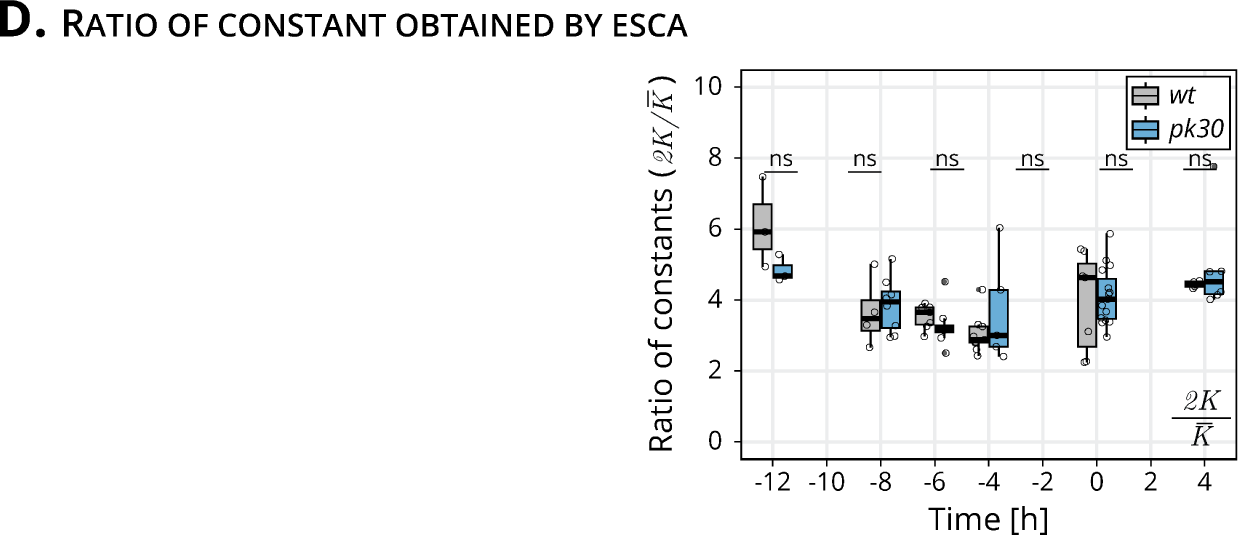
Ratio of elastic constants throughout morphogenesis: (D) Ratio of elastic constants 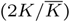 for *wt* and *pk30* (blue) throughout morphogenesis (n⩾3). Significance is estimated using the Kruskal-Wallis test. ns, p-val*>*0.05. The time is relative to the peak of cell elongation.

### 4.4 Supplemental Movie 1

Shown here is an example of a linear laser ablation, cutting 3-4 cells, in wild type (left) or *pk30* pupal wings. The movie goes dark during the ablation itself. Thereafter, the tissue displaces. Anterior is up; proximal is left.

## 5 Materials and Methods

### 5.1 Key resources table

| RESOURCE | SOURCE | IDENTIFIERS |
| --- | --- | --- |
| **Experimental models: Organisms/strains** |  |  |
| *D. melanogaster* wt: *w<sup>-</sup>*; *EcadGFP*;; | Huang et al. (2009) |  |
| *D. melanogaster* *pk30*: *w<sup>-</sup>*; *EcadGFP*, *pk30*;; | Gubb et al. (1999) | BDSC 44229 |
| *D. melanogaster* *stbm6*: *w<sup>-</sup>*; *EcadGFP*, *stbm6*;; | Wolff and Rubin (1998) | BDSC 6918 |
| *D. melanogaster* *fmi*: *w<sup>-</sup>*; *EcadGFP*, *fmi*;; | Wolff and Rubin (1998) | BDSC 6967 |
| **Chemicals** |  |  |
| Euparal | Carl Roth | 7356.1 |
| Holocarbon oil 700 | Sigma-Aldrich | H8898 |
| Isopropanol (2-propanol) | Sigma-Aldrich | 1.01040 |
| **Software and algorithms** |  |  |
| Fiji (v. 2.0.0-rc-68/1.52e) | Schindelin et al. (2012) |  |
| Ilastik (v. 1.2.2) | Berg et al. (2019) |  |
| MATLAB (v. 9.2.0.1226206 (R2017a)) | MATLAB (2017) |  |
| PreMosa | Blasse et al. (2017) |  |
| R (v. 3.4.1) | R Core Team (2020) |  |
| RStudio (v. 3.6.1) | RStudio Team (2020) |  |
| TissueMiner (v. TM_1.0.2) | Etournay et al. (2016) |  |
| **Other equipment** |  |  |
| Coverslip | Paul Marienfeld GmbH | 0107052 |
| Microscope slides | Paul Marienfeld GmbH | 1000200 |
| Dumont #55 Forceps | Fine Science Tools | 11295-51 |
| Vannas Spring Scissors | Fine Science Tools | 15000-08 |

### 5.2 Fly husbandry

Flies were maintained at 25°C and fed with standard fly food containing cornmeal, yeast extract, soy flour, malt, agar, methyl 4-hydroxybenzoate, sugar beet syrup and propionic acid. Flies were kept at 25°C in a 12 h light/dark cycle. Vials were flipped every 2-3 days to maintain a continuous production of pupae and adult flies. All experiments were performed with male flies, since they are slightly smaller and therefore the wings require less tiling on the microscope to be imaged than females.

### 5.3 Long-term timelapse imaging of pupal wing morphogenesis

#### 5.3.1 Acquisition

White male pupae were collected, slightly washed with a wet brush and transferred to a vial with standard food. At 16 hAPF, the pupal case was carefully dissected so that the wing would be accessible. The pupae was then mounted onto a 0.017 mm coverslip on a self-built metal dish with a drop of Holocarbon oil 700 (Classen et al., 2008). Pupal wing morphogenesis was imaged every 5 min for approximately 24 h, as in Etournay et al. (2015).

Two different microscopes were used for acquisition of long-term timelapses. All *wt, pk30* and *stbm6* movies were acquired using a Zeiss spinning disk microscope driven by ZEN 2.6 (blue edition). This microscope consists of a motorized xyz stage, an inverted stand, a Yokogawa CSU-X1 scan head, and a temperature-controlled chamber set to 25°C. The sample was illuminated with a 488 nm laser, and the emission was collected using a 470/40 bandpass filter, through a Zeiss 63x 1.3 W/Glyc LCI Plan-Neofluar objective and a Zeiss AxioCam Monochrome CCD camera with a 2×2 binning. The whole wing was imaged in 24 tiles with an 8% overlap. Each tile consisted of 50-60 stacks with a Z-spacing of 1 *μ*m. The laser power was set to 0.1 mW.

The two *fmi* movies were acquired with an Olympus IX 83 inverted stand driven by the Andor iQ 3.6 software. The microscope is equipped with a motorized xyz stage, a Yokogawa CSU-W1 scan head and an Olympus 60x 1.3 Sil Plan SApo objective. The setup was located inside a temperature-controlled chamber set to 25°C. The sample was illuminated with a 488 nm laser, and the emission was collected using a 525/50 bandpass filter. The whole wing was imaged by tiling with 8 tiles with a 10% overlap. Each tile consisted of 50-60 stacks with a distance of 1 *μ*m between them. The laser power was set to 0.75 mW.

#### 5.3.2 Processing, segmentation, tracking and database generation

Raw stacks were projected, corrected for illumination artifacts, and stitched using PreMosa (Blasse et al., 2017). The stitched images of individual timepoints were cropped to fit the wing size, registered using the Fiji plugin “Descriptor-based series registration (2D/3D + t)” and converted to 8 bit with Fiji (Schindelin et al., 2012). The segmentation was performed with the Fiji plugin TissueAnalyzer (Schindelin et al., 2012; Aigouy et al., 2010; Aigouy et al., 2016). Segmentation errors were identified and manually corrected by looking at the cell divisions and deaths masks.

Subsequent processing and quantifications were performed using TissueMiner Etournay et al. (2016). Before generating the relational database, we rotated the movies so that the angle formed by a manually drawn line connecting the sensory organs would be 0. We manually defined the regions of interest, such as the blade, hinge and the anterior and posterior regions, using the last frame of the movie. Next, we generated the relational database containing information about the cellular dynamics during morphogenesis using TissueMiner (Etournay et al., 2016).

We queried and worked with the data using the Dockerized version of RStudio (Nickoloff, 2016), which loads all packages and functions required to work with TissueMiner. Movies were aligned by the peak of cell elongation by fitting a quadratic function around the cell elongation values 40 frames before and after the absolute maximum of cell elongation in the blade region for each movie. The maximum of this curve was identified and set as the timepoint 0 h.

### 5.4 Adult wing preparation and analysis of wing shape

Adult male flies were fixed in isopropanol for at least 12 h. One wing per fly was dissected in isopropanol, transferred to a microscope slide and covered with 50% euparal in isopropanol. Wings were mounted with 50-70 *μ*L 75% euparal/isopropanol.

*wt, pk30* and *stbm6* wings were imaged using a Zeiss widefield Axioscan Z1 microscope equipped with a Zeiss 10x 0.45 Air objective. *fmi* wings were imaged using a Zeiss widefield Axiovert 200M microscope equipped with a Zeiss 5x 0.15 Plan-Neofluar air Zeiss objective.

Wing blade parameters were quantified using a custom-written Fiji macro (Schindelin et al., 2012). The shape or major-to-minor ratio was calculated using a custom RStudio script (R Core Team, 2020; RStudio Team, 2020).

### 5.5 Quantification of the PD component of cell elongation *Q*_*xx*_

Prior to all laser ablation experiments, we acquired a stack of 50 *μ*m thick that was projected using PreMosa (Blasse et al., 2017). We cropped a region that enclosed the region that was ablated, segmented cells using TissueAnalyzer (Aigouy et al., 2010,1) and generated a relational database with TissueMiner (Etournay et al., 2016).

The definition of cell elongation was first presented in (Aigouy et al., 2010) and it describes the angle and magnitude of the tensor. The cell elongation tensor is given by

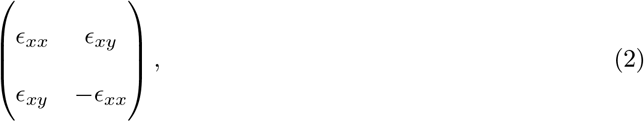

where

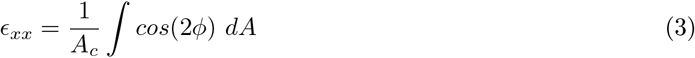

and

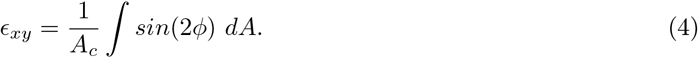

Cell elongation is normalized by the cell area (*A*_*c*_) of each cell. The magnitude of cell elongation is:

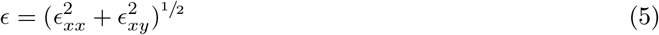

Here we plot *ϵ*_*xx*_ as *Q*_*xx*_, which we describe as the PD component of cell elongation.

### 5.6 Laser ablation experiments

Pupae were dissected and mounted as described for the long-term timelapses. Ablations were always performed in the same region of the wing blade, found in the intervein region between the longitudinal veins L3 and L4 and between the second and third sensory organs. This region was chosen because these landmarks are easily visible in all timepoints. Laser ablations were performed using a Zeiss spinning disc microscope equipped with a CSU-X1 Yokogawa scan head, an EMCCD Andor camera, a Zeiss 63x 1.2 water immersion Korr UV-VIS-IR objective and a custom-built laser ablation system using a 355 nm, 1000 Hz pulsed ultraviolet (UV) laser (Grill et al., 2001; Mayer et al., 2010). The imaging and cutting parameters for line and circular laser ablations are shown in Table 1.

**Table 1:** Parameters used to perform laser ablations.

|  | LINEAR ABLATIONS | CIRCULAR ABLATIONS |
| --- | --- | --- |
| Exposure time [s] | 0.05 | 0.05 |
| 488 nm laser intensity [%] | 50 | 50 |
| Time interval [s] | 0.09 | 2.55 |
| Pulses per shot | 25 | 25 |
| Shots per $\mu\text{m}$ | 2 | 2 |
| Shooting time [s] | 0.67 | 147.28 |
| Thickness of stack ablated [$\mu\text{m}$] | 1 | 20 |

#### 5.6.1 Linear laser ablations to calculate the initial recoil velocity

We performed both types of linear ablations in only one plane of the tissue, in order to minimize the time required for ablation and therefore be able to acquire the initial recoil velocity upon ablation (no imaging is possible during ablation). The length of the linear laser ablations was 10 *μ*m, ablating 3-4 cells. We drew kymographs perpendicularly to the cut to follow the two edges of one ablated cell using Fiji (Schindelin et al., 2012). The initial recoil velocity was calculated as the average displacement of two membranes of the same cell that occurred during the black frames of the ablation itself. This calculation was made using a self-written MATLAB script (MATLAB, 2017). The image acquired prior to the laser ablation was used to compute *Q*_*xx*_ in that region, as described in Subsection 5.5, and the time corresponding to the maximum of cell elongation was defined as 0 h.

#### 5.6.2 Elliptical Shape after Circular Ablation (ESCA)

Circular laser ablations used for ESCA were 20 *μ*m in radius (approximately 10 cells). This radius was selected was that it would fit into the same blade region throughout morphogenesis. Due to the bigger size of these cuts and the curvature of the tissue, we cut the tissue along a stack of 20 *μ*m thick. Approximately 2 min after the ablation, we acquired a stack of 50 *μ*m. This image was projected using PreMosa (Blasse et al., 2017) and preprocessed by applying a Gausian blur (*σ*=1) and background subtraction (rolling ball radius = 30) in Fiji (Schindelin et al., 2012). The next steps were performed as in Dye et al. (2021): the image of the final shape of the cut was segmented using Ilastik (Berg et al., 2019) by defining three regions: membrane, cell and dark regions. The segmented image was thresholded to obtain a binary image of the final shape of the cut. We fitted two ellipses to this image: one to the inner piece and another one to the outer outline of the cut. Based on the shape of these ellipses, the the method outputs the anisotropic 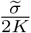 and isotropic stress 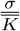 as a function of their respective elastic constants, and the ratio of elastic constants 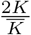. A small number of experiments were fitted poorly (defined as an error per point greater than 0.3) and were therefore excluded from analysis. Prior to the circular ablation, a stack of 50 *μ*m was acquired and used to calculate cell elongation before ablation (Subsection 5.5). The time corresponding to the maximum of cell elongation was set to be 0 h.

#### 5.6.3 Kymograph analysis and fit to model

The ablations used to calculate the mechanical stress along the PD axis for *wt* and *pk30* were further analyzed with the rheological model. To do so, we processed the kymographs by applying a Gaussian blur (*σ*=1) (Schindelin et al., 2012), and then we segmented these kymographs with Ilastik (Berg et al., 2019). Using a self-written Fiji macro (Schindelin et al., 2012), we extracted the intensity profile for each timepoint. Next, we wrote an R script (R Core Team, 2020; RStudio Team, 2020) to identify the membrane displacement over time and obtained a unique curve per kymograph, which could be fitted with our model. We modelled a local patch of tissue as a combination of a spring with spring constant *k*, representing the ablated cells and two KV elements with spring constants *k*_*f*_ and *k*_*s*_ and viscosity coefficients *η*_*f*_ and *η*_*s*_, representing the unablated cells, as shown in (Fig 2C-D). Because the local tissue strain in the experimental measurement is expressed by the displacement of the bond nearest to the ablation, in the rheological model we represent tissue strain by displacements of the two KV elements. In principle, the strain can be recovered by normalising the displacements by the width of ablated cells. Displacements of the two KV elements are defined as a change in the distance between the end points of the KV elements *x*_*i*_(*t*), relative to their initial values *x*_*i*_(0), where *i* ∈ {*f, s*} for fast (*f*) and slow (*s*) element.

Mechanical stress in the tissue is represented by the *σ* acting on our model, and we assume that *σ* is not changed by the ablation. Before the ablation, the model is in mechanical equilibrium and we can write

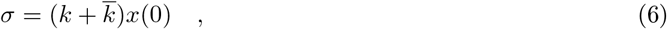

where *x*(0) is the initial distance between the two end points of the model, and 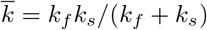 is the elastic constant of the two KV elements connected in series. Upon ablation, the spring *k* is removed and stresses in the model are imbalanced. The distance between the end points of the model *x*(*t*) then evolves towards the new equilibrium position. The distance *x*(*t*) can be decomposed as *x*(*t*) = *x*_*f*_ (*t*) + *x*_*s*_(*t*), where *x*_*f*_ (*t*) and *x*_*s*_(*t*) are the time-dependent distances between end points of the two KV elements, representing their strains. The dynamics of *x*(*t*) is then obtained by writing the force balance equation for the two KV elements

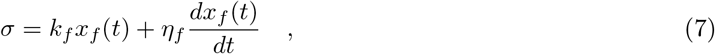

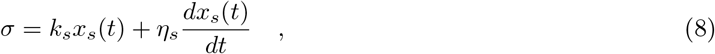

We solve for *x*_*f*_ (*t*) and *x*_*s*_(*t*) to obtain

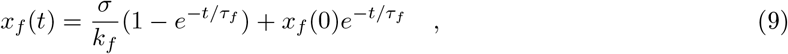

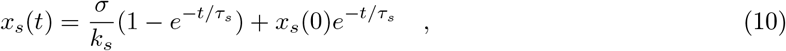

Where

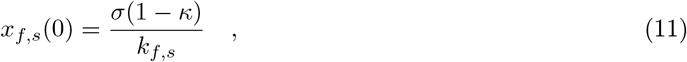

where 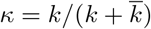is the fraction of the overall model elasticity 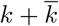 destroyed by the ablation. The displacement relative to the initial configuration Δ*x*(*t*) = *x*(*t*) − *x*(0) is therefore

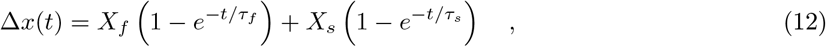

where we introduced the long time displacements associated with the two KV elements

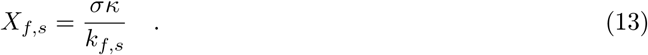

For simplicity, in the main text we refer to the long time displacements *X*_*f*_ and *X*_*s*_ of the two KV elements simply as displacements.

### 5.7 Statistical analysis

Statistical analysis was done using R (R Core Team, 2020; RStudio Team, 2020). We first tested normality of the data using the Shapiro–Wilk test. When data were normal, we used Student’s t-test to test statistical significance between two groups. When data were not normally distributed, significance was tested using the Mann-Whitney U test for two groups or Kruskal-Wallis test for multiple groups. Statistical test results are shown on the figure captions.

## 1 Appendix 1

During the course of this work, we identified a delay on the onset of pupal wing morphogenesis compared to previous work (Etournay et al., 2015). In the past, cells reached their maximum of cell elongation at 22.9±0.4 hAPF, while now they reach it at 28 hAPF (Fig A1.1A). To combine data acquired at different times, we present cell dynamics data aligned in time by the peaks of cell elongation, and we refer to this timepoint as 0 h (Fig A1.1A). We investigated the cell dynamics underlying tissue morphogenesis in the delayed flies and observed that the shear rates were comparable with the older flies (Fig A1.1B). Thus, it is reasonable to shift the curves by aligning them to a new reference time.

**Figure A1.1:**
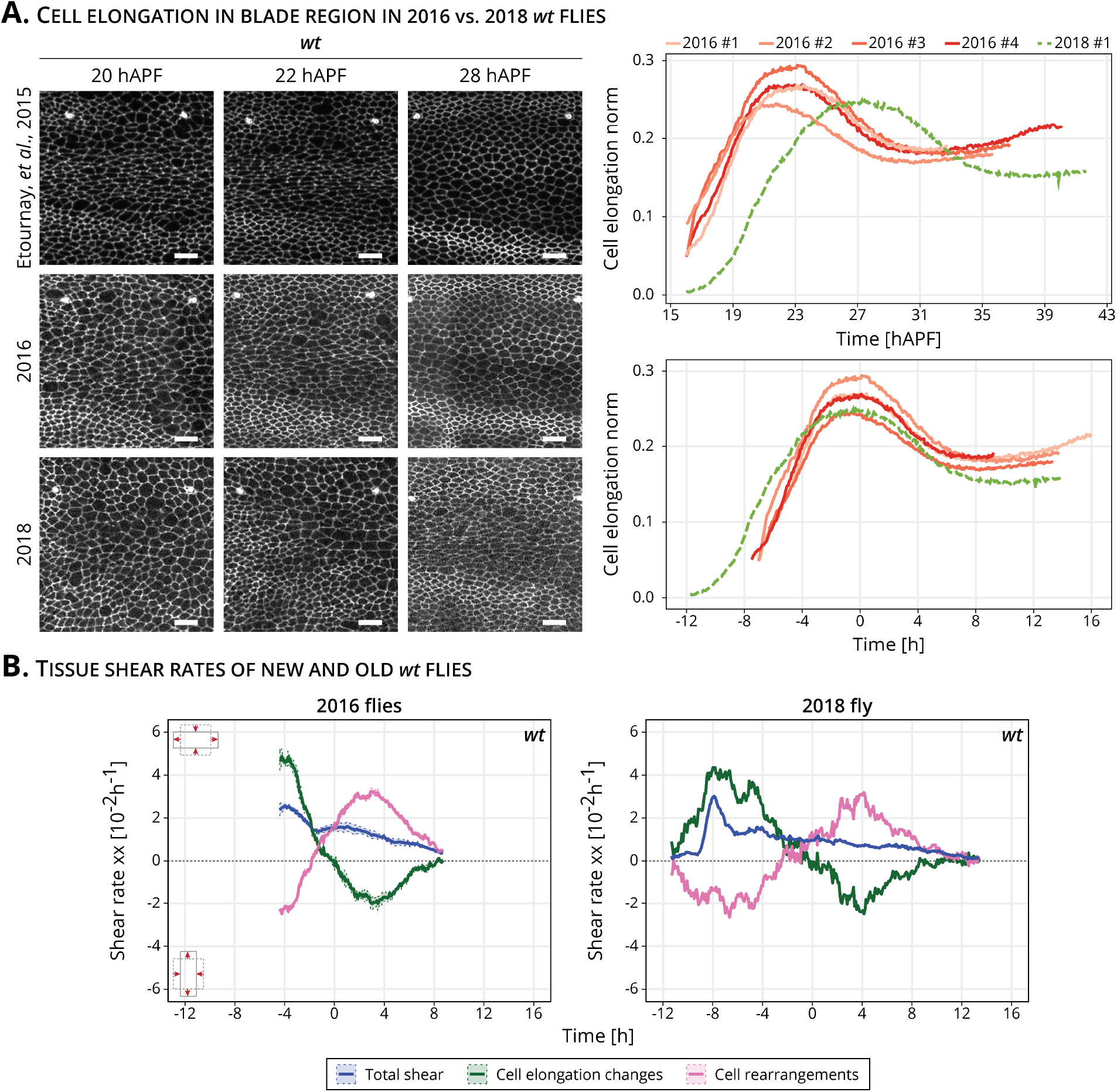
Delay and time alignment of old and newer *wt* flies: (A) Left: snapshots of the blade region of long-term timelapses of pupal wing morphogenesis acquired in different years. Scale bar, 10 *μ*m. Right: cell elongation norm during pupal wing morphogenesis for old flies (orange palette, 2016 flies) and new flies (green curve, 2018 fly). Top plot: cell elongation magnitude for each movie not aligned in time. The peak of cell elongation is delayed from around 23 hAPF to 28 hAPF. Bottom plot: cell elongation magnitude aligned in time. The time 0 h corresponds to the peak of cell elongation. (B) Cell dynamics contributions underlying anisotropic tissue deformation for older (n=4) and newer (n=1) *wt* flies. The time is relative to the peak of cell elongation.

